# Biodiversity impact of the consumption of peat and wood-fired district heating

**DOI:** 10.1101/2024.03.19.585717

**Authors:** Veera Vainio, Sami El Geneidy, Panu Halme, Maiju Peura, Janne S. Kotiaho

## Abstract

The use of biofuels is becoming an increasingly important part of national and corporate climate strategies. At the same time, the consumption-based biodiversity impacts of biofuels are generally poorly known. Here we used a consumption-based approach to assess the biodiversity impacts of peat and wood-fired district heating in Finland. We combined the information on the area of impacted ecosystems and their condition before and after the impact to evaluate the impact as habitat hectares, i.e., the loss in the condition of the impacted habitats. The habitat hectare approach has not been used in previous studies on consumption-based biodiversity impacts but could be replicated to assess biodiversity impacts in different contexts around the globe. We present an eight-step general protocol for such assessment and discuss the usability of the protocol in assessing consumption-based biodiversity impacts of district heating systems. Considering different fuel types, peat had the highest biodiversity impact per unit area, followed by chips from roundwood and logging residue chips. If we consider the impacts per unit energy, chips from roundwood had the highest and peat the lowest biodiversity impact. We conclude that it is possible to assess biodiversity impacts of raw material-based consumption, like we did in our example case. This protocol should be further developed and refined in different systems and with different raw materials.

## 1. Introduction

Energy used in buildings is close to 40 % of the global energy consumption, and heating consumes roughly 60 % of this (Pérez-Lombard et al. 2008). District heating is commonly used in Northern and Eastern Europe as well as in Russia and China (Werner 2017, Paiho and Saastamoinen 2018, Mazhar et al. 2018). For example, about half of the Finnish as well as Swedish population and buildings are dependent on district heating (Eriksson et al. 2007, Paiho and Saastamoinen 2018). Moving away from fossil fuels is likely to increase the demand for wood and other renewable biomass-based district heating (Persson and Werner 2011, Mazhar et al. 2018), and there indeed is some potential in bioenergy (Creutzig et al. 2015, Börjesson et al. 2017). However, expanding the use of bioenergy is already constrained for example by its availability (Szarka et al. 2017, Anttila et al. 2018, Verkerk et al. 2019) and competition of resources between energy and non-energy uses (Mandley et al. 2020). Also, the pressure to increase forest carbon sinks is preventing significant increases in forest biomass harvests (Masson-Delmotte et al. 2018). Biodiversity impacts of the extraction of the biomass is likely to cause yet another constraint (Ranius et al. 2018), but these impacts are still to be quantified in relation to district heating.

Human activity has generated significant changes in the biosphere over the past several decades (Foley et al. 2005, Barnosky et al. 2012, Ellis et al. 2013). The abundance and diversity of nature are declining, species extinction rates are accelerating, and ecosystems are being degraded all over the world (Barnosky et al. 2012, Newbold et al. 2015, IPBES 2018). The most important direct drivers of biodiversity loss and ecosystem degradation are unsustainable land-use and intensive resource use including direct exploitation of species, but also climate change, spreading of harmful alien species, and pollution of habitats (IPBES 2019). It has been estimated that the negative impacts of human land-use already affect up to 75% of the earth’s surface (IPBES 2018). Extracting biomass for district heating can be considered to fall under the category of direct exploitation and it clearly has potential to contribute to biodiversity loss.

Organizations, for example private businesses and public services such as district heating plants, hospitals, and education institutions, are the way we have organized the functions in our societies. To understand the role of organizations in enhancing or reducing planetary well-being (Kortetmäki et al. 2021), we need to be able to identify and quantify the environmental impacts, such as carbon emissions and biodiversity loss their operations are causing. Carbon footprint is a commonly used tool for assessing consumption-based climate impacts of human activities (Wiedmann and Minx 2008), but tools and methodologies to assess consumption-based biodiversity impacts are still less developed but emerging (Lenzen 2014, Marques et al. 2017, Crenna et al. 2020, Lammerant et al. 2021, El Geneidy et al. 2023). So far assessments of biodiversity impacts of any consumption at the level of an organization are rare (El Geneidy et al. 2021, Bull et al. 2022, Taylor et al. 2023).

In this paper we develop a framework for assessing the negative biodiversity impacts of the consumption of peat and wood-fired district heating and apply it to assess the biodiversity impacts of one district heating dependent organization, the University of Jyväskylä.

## 2. Assessing biodiversity impacts

### 2.1 Habitat hectares as a common currency for different biodiversity impacts

In general, human activities can have negative or positive impacts on species or ecosystems. Ecosystem degradation refers specifically to negative or harmful impacts (IPBES 2018). For example, the impact that a construction project causes to any ecosystem, or the harm that logging causes to forest ecosystems can be considered as ecosystem degradation.

Detailed measurement of biodiversity is difficult since even a small area almost anywhere may contain hundreds or thousands of species and even more individuals. In practice it is simply impossible to count them all, and thus biodiversity is not readily amenable to very detailed measurement. In general, biotopes, ecosystems, habitat types or ecological features important for large numbers of species can be categorized and their condition mapped much more readily than occurrences of individual species or their population sizes. Thus, we approach the problem by looking at the condition of the ecosystems rather than species or their population. This approach has also been called the habitat hectares approach (Parkes et al. 2003).

The condition of any ecosystem can be considered to vary from 0 to 1. An area that has been completely degraded obtains value 0, and an area in a completely natural, undisturbed condition obtains value 1. Thus, one habitat hectare equals one hectare of undisturbed ecosystem. For example, a value 0.4 is assigned to an area that has been degraded so that it has lost 60 % of its pristine condition. Habitat hectare is a common currency that can be used when e.g., losses and gains are being balanced in biodiversity offsetting projects (see e.g., Moilanen and Kotiaho 2018, 2020, 2021).

There are also two other methodologies for assessing biodiversity impacts based on consumption: environmentally extended input-output analysis and life cycle assessment (LCA) (Marques et al. 2017, Crenna et al. 2020). While both methods can be used to assess the biodiversity impacts of consumption across the value chain, EEIO analysis is more appropriate for analyzing sectoral impacts, while LCA is more appropriate for analyzing product or process specific impacts (Marques et al. 2017). Here we utilize the habitat hectare approach, which is closer to LCA than EEIO, because we are analyzing the impact of wood and peat for the district heating consumption with process-based impact factors.

The biodiversity impacts of district heating are indirect from the point of view of the organization purchasing the energy. Compared to climate impacts, this would be categorized as scope 2 impact. However, the concrete impact on nature is direct, and could be accounted for as scope 1 impact for the company producing the district heating, or if it is not the same company, for the company actually extracting the wood or peat.

### 2.2 Biodiversity impacts of extracting wood for energy

Forest-based bioenergy can be sourced by extracting additional biomass from production forests or by using industrial by-product residues such as bark and sawdust. Extraction of logging residues results in biomass and thus nutrients removal, and disturbance of soils and several other environmental features important for biodiversity (Ranius et al. 2018). How much biomass is extracted at a site-level varies among countries, with higher values reported from Sweden and Finland than for example from France and North America (Thiffault et al. 2016). Nevertheless, the evidence is clear that logging residue extraction can have significant negative impacts on biodiversity (Ranius et al. 2018).

Dead wood is an important resource for forest biodiversity. Woody biomass removal reduces the food and habitat available for dead wood dependent organisms (Bouget et al. 2012). Energy-wood extraction decreases the number of wood-inhabiting fungi due to decreased number and volume of habitat patches (Toivanen et al. 2012) and affects many other species groups by reducing the availability of shelter and nesting sites (Ecke et al. 2002). For insects the energy-wood extraction means actual removal of a large number of individuals from forest sites, because insects colonize the dead wood piles before the wood is transported to power plants (Hedin et al. 2008). Physical and geochemical changes due to logging residue extraction also affect field layer vegetation. Overall levels of soil organic matter and soil nutrients are reduced, with a potential loss of soil fertility and site productivity. Changes in soil physical properties due to increased heavy machinery traffic in the forest, thinner protective mats of logging residue, and changes in soil moisture and organic matter are examples of the changes caused by the extraction (Verkerk et al. 2019).

### 2.3 Biodiversity impacts of extracting peat for energy

In Finland many peatlands are threatened ecosystems and host a high number of threatened species. Currently the main threat to peatlands is habitat destruction due to ditching, peat mining, forestry, and agriculture (Alanen and Aapala 2015, Hyvärinen et al. 2019, Kontula and Raunio 2019). Peat mining has a detrimental effect on peatlands. Whole ecosystems are destroyed when peatlands are drained, and all the surface vegetation is removed (Finnish Ministry of the Environment 2015). The direct impact of removing the surface layer including all the flora and fauna is evident, but indirect biodiversity impacts outside the direct extraction area via changes in hydrology can also occur. Hydrology is one of the most important features of peatland ecosystems and it largely determines peatland ecosystem functions and species diversity (Konar et al. 2013, Haapalehto et al. 2014, Kareksela et al. 2015).

Drainage of the extraction site unavoidably also affects the hydrology of the surrounding areas by lowering their groundwater levels (Tahvanainen 2011, Alanen and Aapala 2015, Paal et al. 2016). Drainage can also increase nutrient leaching and the flow of suspended solids to downstream water systems (Finnish Ministry of the Environment 2015). These changes degrade the peatland ecosystems (e.g., Mazerolle 2003).

## 3. Framework for assessing the biodiversity impact of district heating consumption of an organization

There are several steps that need to be taken to assess an organization’s biodiversity impact. Here we develop a framework with eight steps as illustrated in **Figure 1**. In this section, we describe each of the steps in detail and apply the framework to the district heating consumed by the University of Jyväskylä.

**Figure 1.**
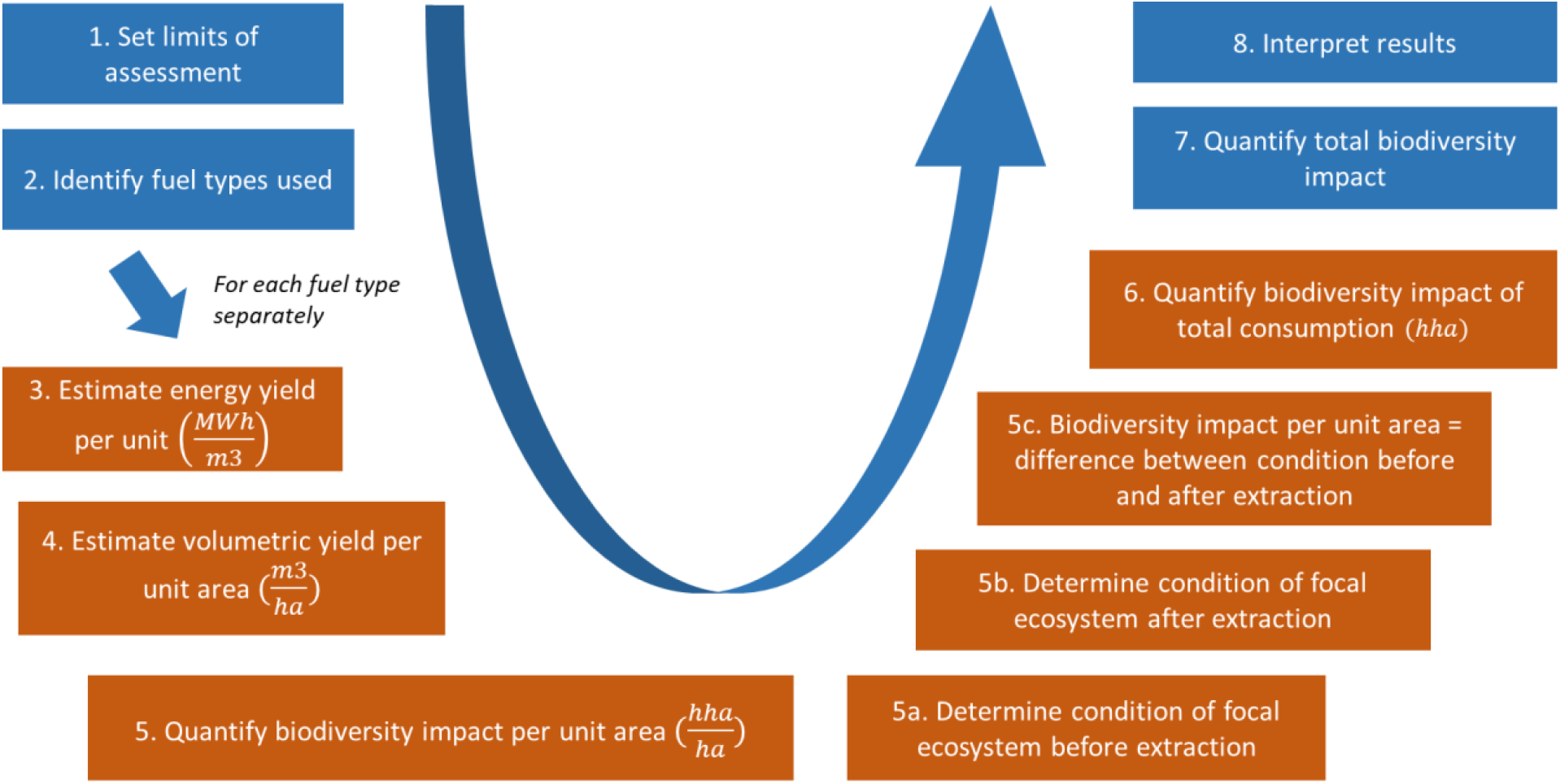
Framework and steps for assessing the biodiversity impact of the consumption of peat and wood-fired district heating of an organization. Blue color indicates common steps for all fuel types and orange color indicates the calculation steps that need to be completed separately for each fuel type.

### Step 1. Set the limits of the assessment

To start the assessment, one needs to delimit the system in focus. In other words, one needs to determine which parts and operations of the organization will be included in the assessment of the biodiversity impacts. Once the limit has been decided, the total heat consumption of the delimited part needs to be calculated.

In the current study we focused on the consumption of district heating of the University of Jyväskylä and thus decided to include buildings used by the University that are located within the City of Jyväskylä district heating network. The areas of the buildings rented out for other organizations were deduced from the total consumption of each of the buildings. Data was available for the years 2019-2021, and 2021 is used as an example in the text. The other two years are included in **Table 1**.

**Table 1.**
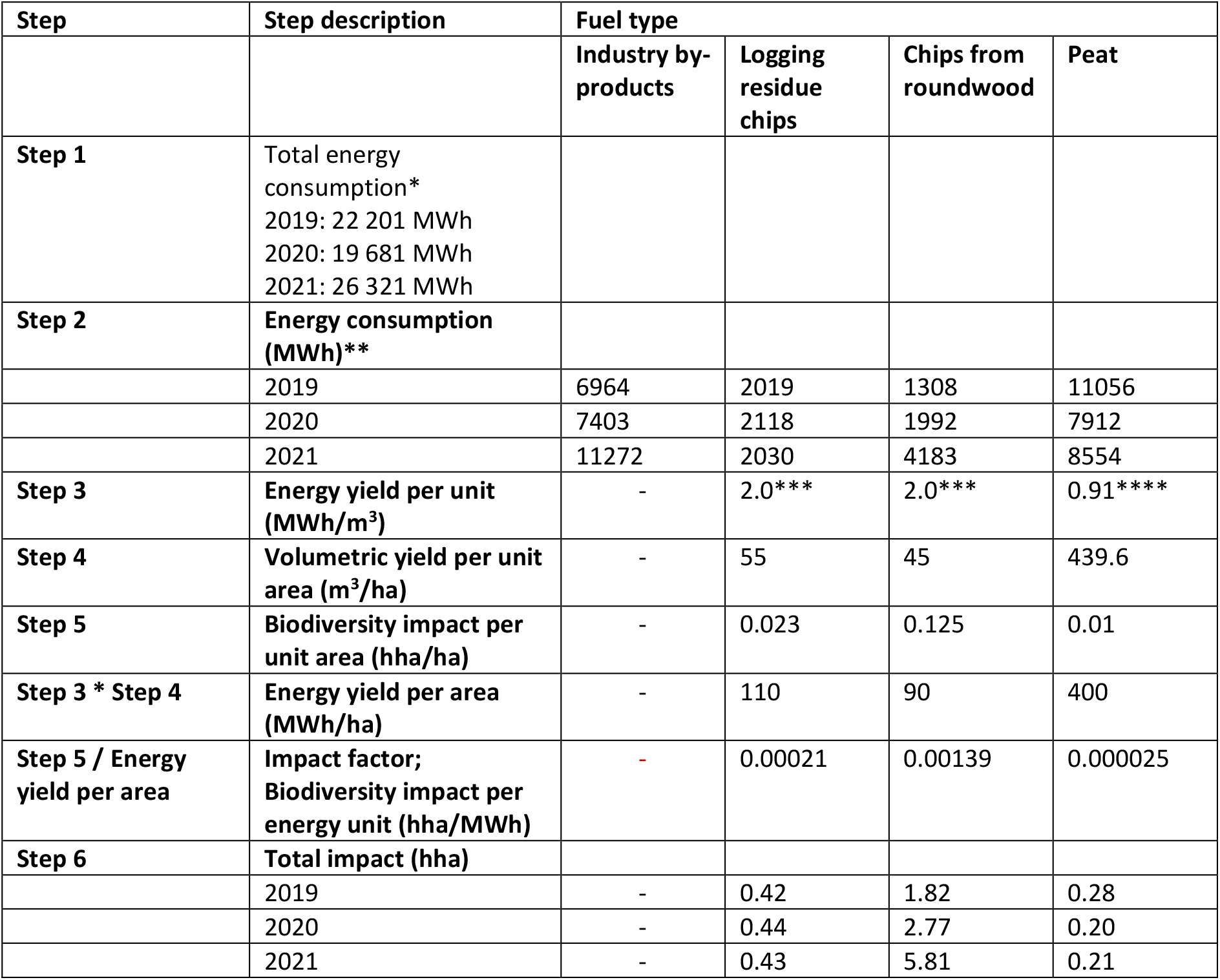
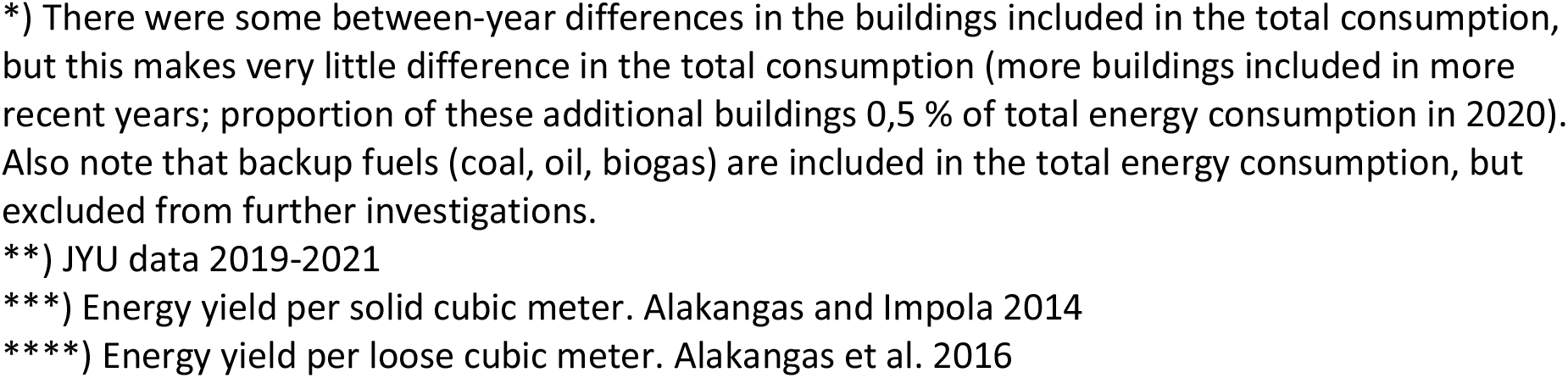

| Step | Step description | Fuel type |  |  |  |
| --- | --- | --- | --- | --- | --- |
|  |  | Industry by-products | Logging residue chips | Chips from roundwood | Peat |
| **Step 1** | Total energy consumption*<br>2019: 22 201 MWh<br>2020: 19 681 MWh<br>2021: 26 321 MWh |  |  |  |  |
| **Step 2** | **Energy consumption (MWh)\*\*** |  |  |  |  |
|  | 2019 | 6964 | 2019 | 1308 | 11056 |
|  | 2020 | 7403 | 2118 | 1992 | 7912 |
|  | 2021 | 11272 | 2030 | 4183 | 8554 |
| **Step 3** | **Energy yield per unit (MWh/m<sup>3</sup>)** | - | 2.0*** | 2.0*** | 0.91**** |
| **Step 4** | **Volumetric yield per unit area (m<sup>3</sup>/ha)** | - | 55 | 45 | 439.6 |
| **Step 5** | **Biodiversity impact per unit area (hha/ha)** | - | 0.023 | 0.125 | 0.01 |
| **Step 3 \* Step 4** | **Energy yield per area (MWh/ha)** | - | 110 | 90 | 400 |
| **Step 5 / Energy yield per area** | **Impact factor; Biodiversity impact per energy unit (hha/MWh)** | - | 0.00021 | 0.00139 | 0.000025 |
| **Step 6** | **Total impact (hha)** |  |  |  |  |
|  | 2019 | - | 0.42 | 1.82 | 0.28 |
|  | 2020 | - | 0.44 | 2.77 | 0.20 |
|  | 2021 | - | 0.43 | 5.81 | 0.21 |

\*) There were some between-year differences in the buildings included in the total consumption, but this makes very little difference in the total consumption (more buildings included in more recent years; proportion of these additional buildings 0,5 % of total energy consumption in 2020). Also note that backup fuels (coal, oil, biogas) are included in the total energy consumption, but excluded from further investigations.
\*\*) JYU data 2019-2021
\*\*\*) Energy yield per solid cubic meter. Alakangas and Impola 2014
\*\*\*\*) Energy yield per loose cubic meter. Alakangas et al. 2016

Note that this study focuses on the direct ecological impacts of extracting fuels for district heat production. Therefore, indirect impacts from e.g., the machinery needed for extraction, or the infrastructure needed for heat production are excluded.

The total consumption of district heating at the University of Jyväskylä was 26 321 MWh in 2021.

### Step 2. Identify the fuel types used

The fuel types and the share of each with which the heat is produced must be identified and an understanding about how and from where the fuels are being extracted needs to be developed.

University of Jyväskylä purchases the district heating from the local energy company Alva that is owned by the City of Jyväskylä. Alva produces around 1 TWh of district heat annually (Alva-yhtiöt Oy 2023).

The power plants of Alva are combined heat and power plants (CHP), so they produce both heat and electricity. We assumed that the total fuel composition of these plants also applies to their district heat production. The energy is produced by burning wood fuels (66.5 % in 2021) and peat (32.5 %). Coal, oil, and biogas are burnt in small quantities (1.0 %) as backup fuels (Alva-yhtiöt Oy 2022, **Figure 2**). We decided to assess the biodiversity impacts of consuming district heat produced using the main fuels, wood and peat, and excluded the imported fossil fuels from the analysis. Wood fuels were comprised of forest fuels and industry residues. Forest fuels are further divided to logging residue chips and to chips from roundwood. Logging residue is mostly extracted from clear-cut sites, whereas roundwood is mostly extracted from young forest stands.

**Figure 2.**
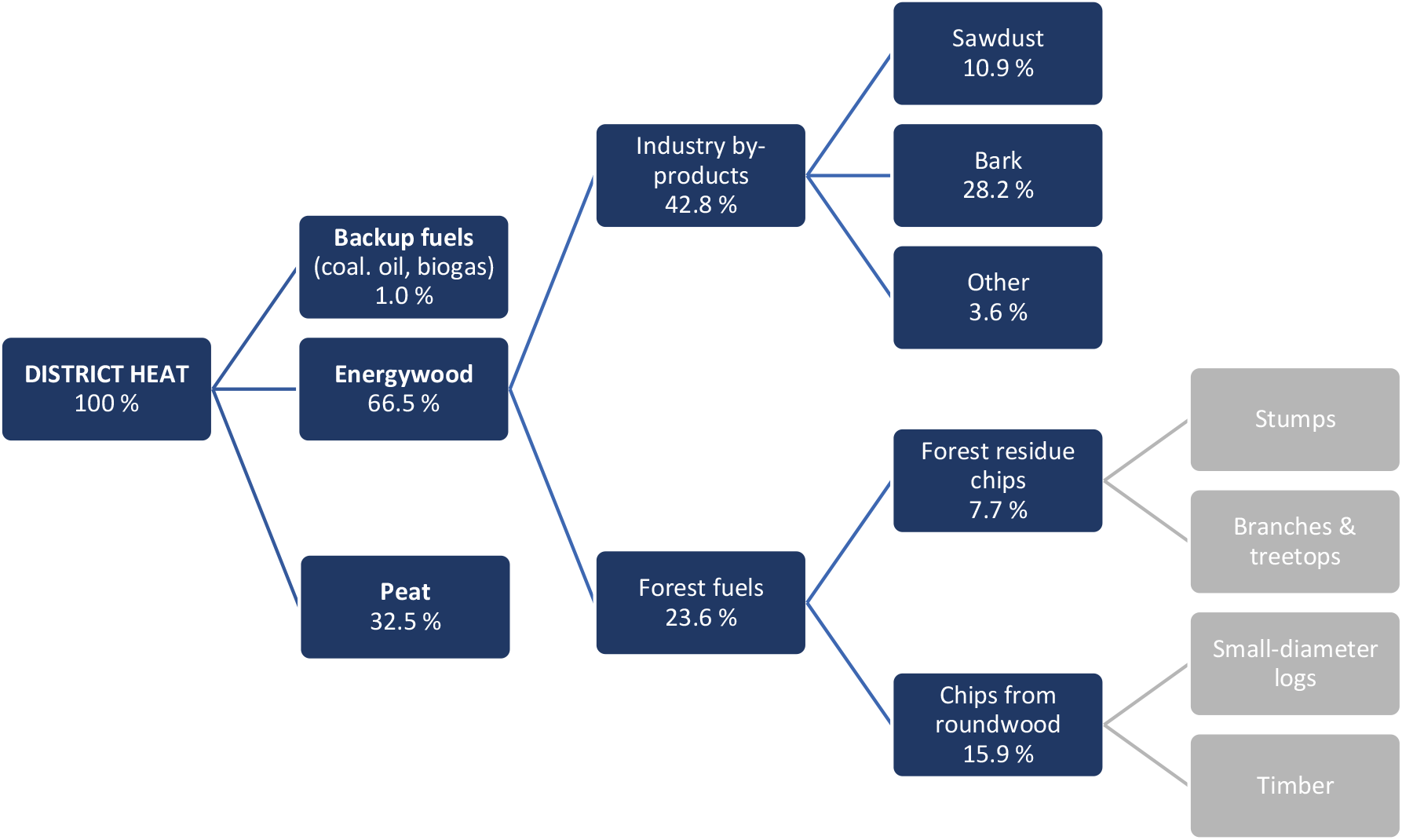
The fuel types and the share (percentage of the total energy produced) of each that are used for district heat production in the City of Jyväskylä (first level Alva-yhtiöt Oy 2022, further classifications OSF 2021). Note that stumps are not used by Alva and thus their extraction is not part of this study.

**Figure 2.**
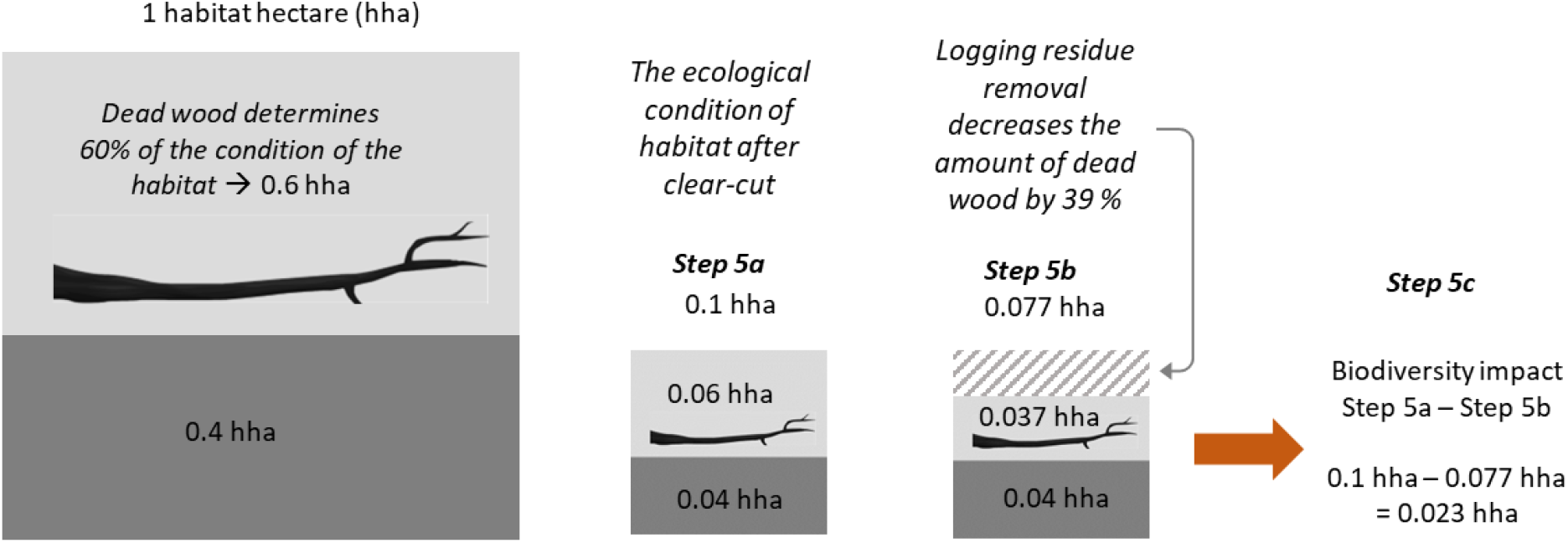
Quantifying the biodiversity impact per unit area for logging residue chips. The assessed impact is logging residue removal from clear-cuts. Before the extraction the ecological condition of focal ecosystem, i.e., clear-cut area, is 0.1 habitat hectares and dead wood determines 60 % of the condition (Step 5a). Logging residue removal decreases the amount of dead wood in clear-cut area by 39 %, and thus the ecological condition of focal ecosystem after impact is 0.077 habitat hectares (Step 5b). Biodiversity impact per unit area is the difference in condition before and after the impact which is 0.023 hha/ha (Step 5c).

**Figure 3.**
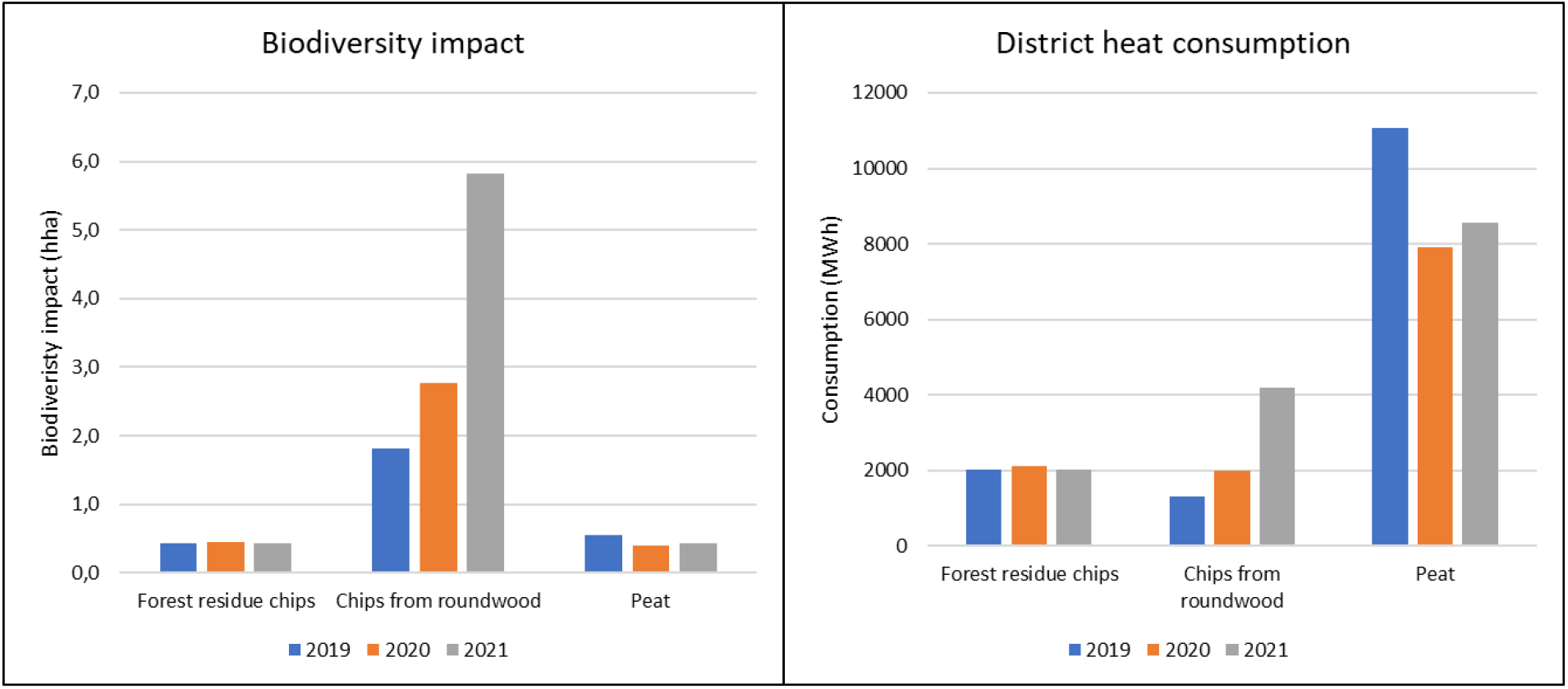
The biodiversity impact (hha) of district heating use by fuel type and the consumption of each fuel type (MWh) in the University of Jyväskylä in 2019-2021.

Residues from timber industry include bark chips, wood residue chips and sawdust. Logging is necessary for obtaining industrial wood residue, but the residues are not the primary reason for logging. However, it is worth noting that if an increase in the demand for timber industry wood residue would encourage more logging, then the use of residues in district heating would cause an additional biodiversity impact. The demand and price of timber industry wood residue is currently low compared to timber wood (Metsälehti 2023). Therefore, we considered that the biodiversity impacts of industrial wood residue have zero biodiversity impact in the case of district heating users. The biodiversity impact of logging for timber should be addressed to the primary sawmill industry.

Peat was assumed to be milled peat, which is the main form of peat used in district heating systems in Finland (Alakangas et al. 2016).

Of the total consumption of the University of Jyväskylä, 11 272 MWh was produced from timber industry residues, 2 030 MWh from logging residue chips, 4 183 MWh from roundwood chips, and 8 554 MWh from peat in 2021 (**Table 1**).

### Step 3. Estimate the energy yield per volumetric unit of each fuel type

#### Steps 3-6 need to be conducted separately for each of the fuel types

The energy yield per volumetric unit of each fuel type (MWh/m^3^) can be derived from literature. We utilized national statistics from the Technical Research Centre of Finland (Alakangas and Impola 2014, Alakangas et al. 2016). The energy yields per volumetric unit for the different fuel types included in the assessment are listed in **Table 1**.

### Step 4. Estimate the volumetric yield of each fuel type per unit of extraction area

The volumetric yield per unit of extraction area can be estimated based on literature. Detailed information on the different extraction practices of the fuel types is also necessary. This process is next described separately for each fuel type. For timber industry residues this step is not necessary, as we assumed that the negative biodiversity impact due to logging for timber should be addressed for the timber sawmill industry and not for the district heating industry using their residues. It should be noted that the volumetric yield varies by region and country and each assessment should be based on the local context.

#### Logging residue chips

Logging residue contains branches and treetops from commercial logging, and it is customarily collected only after final felling and not from thinnings (Äijälä et al. 2014, Koistinen et al. 2019). Therefore, we assumed that all logging residue originated from final felling sites. The main method for final felling in Central Finland is clear-cutting (OSF 2018). The extraction of logging residues after clear-cutting causes an additional biodiversity impact because the amount of decaying wood material is significantly reduced (Mikkonen et al. 2018).

The average yield of logging residue is 50-60 m^3^/ha in Finland (Viitasaari 2013). Assuming that on average 55 m^3^ of logging residue is extracted from an average hectare of clear-cut site, to satisfy the consumption of 2030 MWh of heat, when the energy yield of logging residues is 2 MWh/m^3^ (see **Table 1**, Step 1), a total of 18.4 hectares of clear-cut site needs to be harvested for logging residues. Later in the calculations we use the heat produced per hectare which can be obtained by multiplying the amount of logging residue extracted from the average hectare (55 m^3^) by the energy yield of logging residue per volume (2 MWh/m^3^), and that is 110 MWh/ha (see **Table 1**).

#### Chips from roundwood

Roundwood from first thinning of young stands, where the average chest-height diameter is 8-16 cm, is used for energy production (Äijälä et al. 2014). Thus, the assessed biodiversity impact is the thinning of young stands.

The yield of timber from the first thinning is commonly 30 – 60 m^3^/ha (Asikainen et al. 2012, Lauhanen et al. 2014, Karttunen et al. 2016). Assuming that on average 45 m^3^ of roundwood is extracted from an average hectare of thinning of young stands, to satisfy the consumption of 4183 MWh of heat, when the energy yield of roundwood chips is 2 MWh/m^3^ (see **Table 1**), a total of 46.5 hectares of thinning needs to be conducted. By multiplying the amount of roundwood extracted from the average hectare (45 m^3^) by the energy yield of roundwood per volume (2 MWh/m^3^) we get the energy yield per unit area, which is 90 MWh / ha (see **Table 1**).

#### Peat

Energy yields per hectare were readily available for peat in the literature. To follow the same steps as with other fuel types, the volumetric yield per hectare was calculated by dividing the energy yield per hectare by the energy yield per volumetric unit of peat (0.91 MWh/m^3^, see **Table 1**).

The average yields of energy peat in all of Finland vary from 400 to 500 MWh/ha (Väyrynen et al. 2008). A significant proportion of the peat utilized is produced outside Central Finland (Flyktman 2012) but detailed information on the origin of the peat is not available. Therefore, we assumed that all peat used in the energy production of the local energy company would be produced in Central Finland. The average annual production rate of the peat production sites in Central Finland is 400 MWh worth of energy per hectare (Flyktman 2012). Thus, the volumetric yield of peat is 439.6 m^3^/ha (400 MWh/ha divided by 0.91 MWh/m^3^).

When on average 439.6 m^3^ of peat is extracted per hectare, to satisfy the consumption of 8554 MWh of heat from peat, when the energy yield of peat is 0.91 MWh/m^3^ (see Table 1), a total of 21.4 hectares of peat mining is needed. y multiplying the amount of peat extracted from the average hectare (439.6 m^3^) by the energy yield of peat (0.91 MWh/m^3^) we get the energy yield per unit area, which is 400 MWh/ha (see table 1).

### Step 5. Quantify the biodiversity impact per unit area for each fuel type

To assess the biodiversity impact of the extraction of each of the fuel types in habitat hectares, information on three aspects of each of the ecosystems where extraction is performed is needed: i) the ecosystem condition of the extraction area prior to the extraction (habitat hectares per hectare (hha/ha)), ii) the ecosystem condition of the extraction area after the extraction (hha/ha), and iii) the size of the area extracted (ha). The biodiversity impact is then calculated as the difference in per hectare condition before and after the extraction. Sometimes several calculations per fuel type may be needed if the ecosystem condition before extraction varies from site to site.

#### Logging residue chips

##### 5a) Determine the condition of the focal ecosystem before the extraction per unit area

Logging residue chips are obtained from biomass that is left behind from forest clear-cutting for other than district heating purposes. In other words, the clear-cutting was not conducted to obtain logging residues for district heating. Thus, in this case the impact of clear-cutting can be excluded from the biodiversity impact calculation and the focus must be on what is the additional biodiversity impact caused by the logging residue extraction.

The impact of forest management actions on ecosystem quality has been assessed by Mikkonen et al. (2018), Moilanen and Kotiaho (2020), and Jalkanen et al. (2023). Based on these studies, we assumed the ecological condition of a clear-cut forest site to be 0.1 habitat hectares per hectare before the extraction of the logging residue.

##### 5b) Determine the condition of the focal ecosystem after the extraction per unit area

When logging residue is removed from the final felling site, the amount of decaying wood is reduced by 39 % (Eräjää et al. 2010). Elsewhere it has been estimated that 60 % of the area’s ecological value is determined by dead wood (Kotiaho et al. 2015, 2016). Thus, by assuming that dead wood comprises 60 % of the 0.1 habitat hectares per hectare of the clear-cut site, the condition of the clear-cut site after the logging residue extraction can be calculated as 0.10 hha/ha - (0.10 hha/ha × 0.60 × 0.39) = 0.077 hha/ha.

##### 5c) Calculate the biodiversity impact per unit area by taking the difference between the condition before and after the extraction

The negative biodiversity impact of logging residue extraction per hectare is 0.1 hha/ha – 0.077 hha/ha = 0.023 hha/ha. The steps 5a to 5c are illustrated in Figure 2.

#### Chips from roundwood

##### 5a) Determine the condition of the focal ecosystem before the extraction per unit area

The condition of the young stand before the thinning has been estimated to be 0.25 habitat hectares per hectare (Moilanen and Kotiaho 2020).

##### 5b) Determine the condition of the focal ecosystem after the extraction per unit area

According to Mikkonen et al. (2018), first thinning reduces the ecological condition of the site by 50 %. Thus, after the first thinning, the condition of ecosystem is 0.50 × 0.25 hha = 0.125 habitat hectares per hectare.

##### 5c) Calculate the biodiversity impact per unit area by taking the difference between the condition before and after the extraction

The biodiversity impact of first thinning equals 0.25 hha – 0.125 hha = 0.125 habitat hectares per hectare of thinned area.

#### Peat

The calculation of the biodiversity impact of peat as a fuel for district heating is very different from that of wood fuels. To obtain the peat, the biodiversity impact occurs once when the peatland is ditched and mining is started, but the same peat extraction site can be used for approximately 15-30 years (Väyrynen et al. 2008). In other words, the fuel is extracted and thus heat obtained for a long period of time from the same site, and when the biodiversity impact of the peat-fired district heating is calculated the duration of mining needs to be taken into account.

Draining the peatland by ditching and removing the top layer of soil with all the vegetation while preparing the site for production causes the most significant impacts of peat mining (Finnish Ministry of the Environment 2015). Therefore, the biodiversity impacts do not significantly increase after initial draining and removal of vegetation.

However, draining of peat extraction sites also affects the hydrology of surrounding peatlands (Finnish Ministry of the Environment 2015). The extent of the draining effect depends on features like topography, peat layer depth, and drainage ditch depth (Paal et al. 2016). Moreover, a common requirement for peat extraction in Finland is that the peat layer is at least 1.5 meters thick. This excludes e.g., the edges of the peatland from peat production. Nevertheless, ditching does drain these areas of the peatland as well, so this area in addition to the actual production area is degraded (Kareksela et al. 2013).

It is not exactly known how large proportion of each drained peatland is suitable for peat mining. We assumed that 1/2 of the drained area is either too shallow or otherwise unsuitable for peat mining (assumption is derived from the data in Kareksela et al. 2013). This means that if one hectare of peatland is being used for peat extraction, another hectare of non-mineable peatland also gets degraded. Thus, the biodiversity impact caused by peat mining should include the impact of peat extraction as well as the impact of draining the non-mineable areas.

##### 5a) Determine the condition of the focal ecosystem before the extraction per unit area

Since no pristine peatlands are taken into use for peat production (Ministry of Agriculture and Forestry of Finland 2012), we assumed the condition of a peatland before preparations for peat extraction to be 30 % of its natural state, or 0.3 habitat hectares per hectare. This is the average condition reported for the drained peatlands suitable for wood production or peat mining in Finland (Kotiaho et al. 2015).

The ecological condition of the non-mineable area is similar to the mineable area before peat extraction, i.e., 0.3 habitat hectares per hectare. Note that if the area would be in a better condition before the extraction, the negative biodiversity impact would be higher that what is calculated below.

##### 5b) Determine the condition of the focal ecosystem after the extraction per unit area

The ecological condition of a peat extraction site is only 0.01 habitat hectares per hectare (Kotiaho et al. 2015). Since peat is not extracted from the non-mineable parts of the peatland, the ecological impact in non-mineable parts is likely to be less severe. We assumed the condition of non-mineable parts to decline a further 50 % due to the draining. The ecological condition of drained non-mineable site is thus 0.3 hha/ha × 0.5 = 0.15 habitat hectares per hectare.

##### 5c) Calculate the biodiversity impact per unit area by taking the difference between the condition before and after the extraction

The ecological condition of the peat extraction site declines from 0.3 hha/ha before peat extraction to 0.01 hha/ha after extraction, and the impact is 0.3 - 0.01 = 0.29 habitat hectares per hectare of extraction site. In non-minable parts, the biodiversity impact of draining is 0.3 - 0.15 = 0.15 habitat hectares per hectare. In reality it is likely that the more intense drainage degrades the area over some years, but to aid the calculation we make a simplifying assumption that the biodiversity impact of draining is full from the beginning of the peat extraction. Because one non-mineable hectare gets drained per each mined hectare, we need to take the average over the two hectares to arrive to the biodiversity impact of mining in habitat hectares per hectare: (0.29 hha/ha + 0.15 hha/ha) ÷ 2 = 0.22 hha/ha.

As the peat is extracted and thus heat obtained for a long period of time from the same site with no additional biodiversity impacts, the biodiversity impacts of peat production should be divided by the peat production years. District heat is consumed in different quantities by many organizations every year, and the division is needed to allow the partitioning of the annual biodiversity impact among the organizations based on their annual consumption. The total biodiversity impact per extraction area hectare divided for the production years of the site (22.5 years on average, Väyrynen et al. 2008) equals 0.22 hha/ha ÷ 22.5 years = 0.010 habitat hectares per year per extraction area hectare.

### Step 6. Quantify the total biodiversity impact of each fuel type consumption

Now that we know the total consumption of the heat at the focal organization (Step 1), fuel types and the share of each with which the heat is produced (Step 2), the energy yield per volumetric unit of each fuel type (Step 3), the volumetric yield of each fuel type per unit area extracted (Step 4), and the biodiversity impact of each fuel type per unit area extracted (Step 5), we can quantify the total biodiversity impact of each fuel type.

To achieve the energy yield per extracted area, the energy yield per volumetric unit (MWh/m^3^) of each fuel type from Step 3 is multiplied by the volumetric yield of each fuel type per unit area extracted (m^3^/ha) from Step 4 (**Eq. 1**). To achieve the biodiversity impact per energy unit (hha/MWh) for each fuel type, the biodiversity impact per unit area extracted (hha/ha) obtained in Step 5 is divided by the energy yield per area extracted (MWh/ha, obtained with Eq. 1) (**Eq.2**). To achieve the biodiversity impact of energy consumption of each fuel type, biodiversity impact per energy unit (hha/MWh) is multiplied by the total consumption of energy produced with that fuel type (**Eq. 3**). Note that these calculations are done separately for each fuel type *i*. The results are summarized in Table 1.

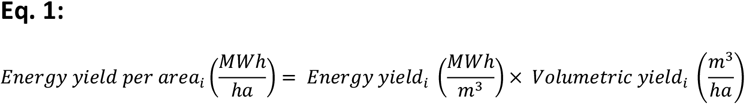

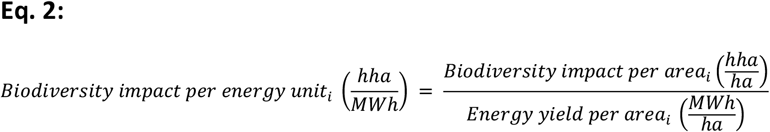

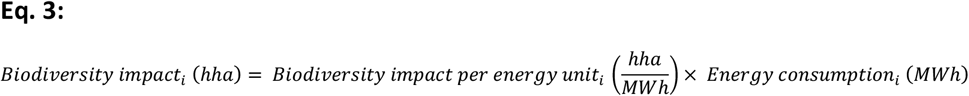

In **Equation 4** we provide the biodiversity impact of energy consumption of fuel type *i* directly without intermediate steps from equations 1, 2 and 3.

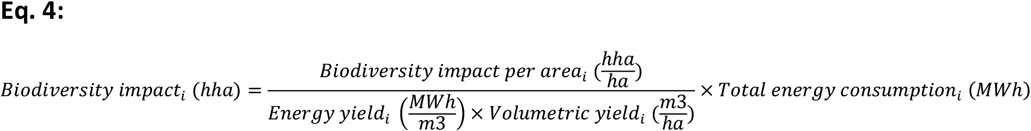

The last two steps of the framework combine all different fuel types.

### Step 7. Quantify the total biodiversity impact of district heating

To illustrate the total biodiversity impact, the impacts of the same ecosystem type can be added together because the biodiversity impact is being measured as the overall impact on the ecosystem with a unified metric: habitat hectare. Adding impacts over the ecosystem types is also possible to illustrate the overall impact. However, this should be done with caution to avoid e.g., fading out the potentially small effects on especially vulnerable ecosystems. The biodiversity impact of district heating consumption of the University of Jyväskylä in 2021 was 6.24 habitat hectares in forest ecosystems and 0.21 habitat hectares in peatlands. This means that ecological condition equivalent to 6.24 hectares of pristine forest and 0.21 hectares of pristine peatland per year is lost due to district heating consumption of the University.

### Step 8. Interpret the results

When interpreting the results, it is important to analyse both the total biodiversity impact of each fuel type, and the biodiversity impact of each fuel type per energy unit (impact factor). The total biodiversity impact indicates the actual impact of the organization on biodiversity and can be used to e.g., formulate mitigation strategies. However, the impact factors are also important, as they can be used to compare different energy sources to choose more sustainable alternatives.

The results can also be used for evaluating biodiversity offsetting actions to reach no-net-loss (NNL) or net positive impact (NPI) for biodiversity. In some cases, it might be beneficial to relate the results to the size of the organization, for example report results as biodiversity impact per employee or per unit area of premises.

It is also important to scrutinize the assumptions behind the calculations. For example, here we focused only on the biodiversity impacts from the land use and ignored the impacts that are due to e.g., emissions from the heat production and their effects on climate change. These and other matters will be further discussed from the viewpoint of our case study in the following section.

## 4. Conclusions

In this paper we presented a general protocol for assessing the biodiversity impact of peat- and wood-fired district heating. With adequate data, the protocol can be used to assess the biodiversity impact of any biofuel-based district heating system, or in fact any other industry where biomass is extracted and utilized. We adopted the habitat hectare approach to make a holistic biodiversity impact assessment when the condition of the focal ecosystem is known before and after the extraction of the biomass. The framework proposed in this paper can be used anywhere in the world, but it is crucial that all the factors and details are adapted to the local context. This includes finding out all the necessary details of local energy production, as well as where and how the different fuels are extracted, by which methods, and what is the ecological impact caused by these practices in the location in question.

Of the fuel types analyzed in this study, peat had the most intensive biodiversity impact per unit area (0.22 hha/ha), followed by roundwood chips (0.125 hha/ha) and logging residue chips (0.023 hha/ha). However, if we consider that peat is extracted annually from the same site with no additional negative biodiversity impacts, the average impact per hectare per annum decreases to 0.01 hha/ha. When we are looking at the biodiversity impact per energy unit, which may in fact be more relevant in the case of energy consumption, the order of the biodiversity impacts is the opposite: largest biodiversity impact (0.00139 hha/MWh) comes from firing roundwood chips, followed by firing logging residue chips (0.00021 hha/MWh), and firing peat had the least biodiversity impact at 0.000025 hha/MWh. Roundwood chips have 56 times greater and logging residue chips 8 times greater biodiversity impact per unit of energy than peat.

To better understand the magnitude of the biodiversity impacts of district heating, we scaled the results to assess the biodiversity impact of total wood- and peat-fired district heating consumption in Finland. In 2021 a total of 40 798 000 MWh of district heating was produced in Finland, of which 42.1 % was from wood fuels and 9.5 % from peat fuels (Statistics Finland 2023). When we apply the biodiversity impact factors derived in this study (see **Table 1**) by taking the average biodiversity impact factor for wood fuels (0.0008 hha/MWh) and utilizing directly the 0.000025 hha/MWh factor for peat fuels, the biodiversity impact would be around 13 700 habitat hectares in forests and 100 habitat hectares in peatlands. For comparison, on average 6500 hectares of forest is voluntarily protected in Finland annually (Koskela et al. 2020). In conclusion it seems that the biodiversity impact of district heating in Finland is remarkable.

This study also provides interesting insights for the political discussion between peat and wood-fired heating in Finland and Europe, which has been further fueled by the ongoing energy crisis in Europe. As a part of their climate targets, the Finnish Government (2019) has been planning to halve the energy use of peat by 2030. Peat burning comprised of 18.9 % of total district heating production in Finland in 2011 and 9.5 % in 2021, indicating a steady decrease (Statistics Finland 2023). In fact, peat production has dropped even faster than expected (Yle News 2022). However, at the same time there has been a steady increase in the use of wood fuels in Finnish district heating, climbing up from 22.3 % in 2011 to 42.1 % of total district heating production in 2021 (Statistics Finland 2023). In the light of our results, this clear shift from peat to wood firing is likely to cause a significant increase in the negative biodiversity impacts of district heating. The trade-off between the climate and biodiversity impacts is truly substantial.

Peat is considered to be a non-renewable fuel and as such especially detrimental to climate like any other fossil fuel (Horsburgh et al. 2022). In contrast, wood fuels are classified as renewable and carbon neutral, even though burning as such is by no means carbon neutral (Schultze et al. 2012). Burning wood is classified carbon neutral because the emissions from wood fuels are allocated to the land use sector at the point of wood extraction and the emissions are assumed to be sequestered by regrowth of the renewable biomass. The classification of wood fuels as carbon neutral has been contested by scientists as it provides no incentive to avoid burning the wood (Norton et al. 2019). In our attempts to reach the carbon neutrality targets, the classification of wood fuels as carbon neutral has given rise to strategies where peat and other non-renewables are replaced by wood fuels. For example, the district heating company Alva, the target of this study, now provides a product called “green heat” that is deemed environmentally friendly. This product consists mostly of wood fuels (Alva 2020), which our results here show are far from environmentally friendly. In addition to climate impacts, future policies and sustainability strategies of energy producers should focus on biodiversity impacts of energy production methods. It is already well recognized that climate change and biodiversity loss require joint solutions and seeking for synergies and avoiding trade-offs (Pörtner et al. 2021), such as revealed by our results. One of the most influential solutions would naturally be to reduce the consumption of energy.

We focused here on land use impacts on biodiversity as it is one of the drivers, if not the key driver for biodiversity loss (IPBES 2019). However, biodiversity loss is also driven by water use, overexploitation of natural resources, climate change, pollution, and invasive alien species (IPBES 2019) and in the future these should also be considered when determining the most sustainable practices of energy production. Relevant for the current study is that it is clear that for both peat mining (Rodhe and Svensson 1995, Kløve 2001) and forest use (IPCC 2019), other significant effects, such as climate change and pollution exist in addition to the biodiversity impacts caused by the fuel extraction processes.

District heating is a necessary part of infrastructure, and it will continue to be used in the future. However, it is important to consider the biodiversity impacts when planning and using district heating. In order to find the least harmful fuel composition for district heating, more research on the biodiversity impacts of different fuels as well as other energy production systems, including non-combustion energy production, are needed.

## Acknowledgements

This work was partly funded by the Strategic Research Council at the Academy of Finland (Kotiaho 345267). We would also like to thank Ulla Helimo and the Sustainability for JYU project members for their support and comments.

## Author contributions

Veera Vainio: Writing – Original Draft, Conceptualization, Methodology, Formal analysis, Investigation, Data curation, Visualization, Project administration

Sami El Geneidy: Writing – Original Draft, Conceptualization, Methodology, Writing – Review & Editing, Supervision

Panu Halme: Writing – Original Draft, Conceptualization, Methodology, Writing – Review & Editing, Supervision, Visualization

Maiju Peura: Writing – Review & Editing, Methodology, Visualization

Janne Kotiaho: Writing – Original Draft, Conceptualization, Methodology, Writing – Review & Editing, Supervision

